# Temporal structure of two call types produced by competing male cicadas

**DOI:** 10.1101/2022.08.14.502195

**Authors:** Takahiro Ishimaru, Ikkyu Aihara

## Abstract

Male cicadas vocalize sounds to attract conspecific females. The acoustic traits of calls vary significantly among species and show unique temporal and spectral patterns that dynamically change, even in the same bout. While the calling behavior of a single cicada has been quantified for many species, the acoustic interaction between multiple cicadas and the usage of different call types have not been well studied. In this study, we examined the interaction between male cicadas (*Meimuna opalifera*) that utilize two types of calls. First, we caught two cicadas in their natural habitat and recorded their calls in the laboratory. Second, we detected the calls of each cicada and classified them into two types: Type I calls with a short duration and high repetition rate and Type II calls with a longer duration and low repetition rate. The analysis of the chorus structure demonstrated that the cicadas vocalized a Type II call immediately after another cicada vocalized a Type I call. Furthermore, we tested the hypothesis that such a timing strategy allowed the cicadas to effectively mask the calls of their competitors. Specifically, we conducted a numerical simulation randomizing the onsets of calls and compared the masking performance with empirical data, which did not support our hypothesis. This study highlights the well-organized structure of cicada calls, even in the choruses with multiple call types, and indicates these calls have a function other than male-male acoustic interaction that requires further investigation.

**Summary statement:** Male cicadas (*Meimuna opalifera*) produce two types of calls by synchronizing their temporal structure and switching call types when positioned close together.

## 1. Introduction

Animals use sounds for various purposes. For instance, frogs and insects form a lek in which males produce sounds to attract conspecific females. Male frogs tend to avoid call overlaps with neighbors (Aihara et al., 2011; Aihara et al., 2014; Brush and Narins, 1989; Gerhardt and Huber, 2002, Jones et al., 2014, Wells, 2007,), whereas insects often overlap their calls (Gerhardt and Huber, 2002; Greenfield and Roizen, 1993; Greenfield et al., 2021). As other examples, male birds sing complex songs for mate attraction (Catchpole and Slater, 2008); monkeys and meercats produce sounds for communication while avoiding call overlaps with each other (Takahashi et al., 2013, Demartsev et al., 2018). Thus, the timing strategy and also the acoustic traits vary significantly among species, playing a vital role for their communication. However, the calling behavior of certain animals has not been well examined due to the lack of high-definition audio recordings. The application of advanced recording techniques, such as microphone array systems (Au and Herzing, 2003; Fujioka et al., 2016; Rhinehart et al., 2020; Suzuki et al., 2018) and the data analysis for the interaction mechanisms (Greenfield et al. 2021, Ota et al., 2020), will contribute to further understanding of animal acoustic communications.

Male cicadas vocalize sounds mainly on tree trunks during the day to attract females. The acoustic traits of calls differ among species (Fonseca, 2014), allowing males to effectively attract conspecific females under acoustically complex conditions including environmental noise and the calls of other animals. Male cicadas have evolved sound-producing organs in their abdomens and can vocalize loud sounds (e.g., up to 148.5 dB sound pressure level in *Cyclochila australasiae*) with melodically complex acoustic traits (Young and Bennet-Clark,1995). In general, many cicadas aggregate at breeding sites and vocalize together, which is called “a chorus center” (Williams and Smith, 1991). The chorus center likely plays a role in attracting females through louder sounds while reducing the predation risk (Hayashi and Saisho, 2015). Cicadas have three ocelli and two compound eyes for perceiving brightness and objects, respectively (Ribi and Zeil, 2015). However, the compound eyes of insects generally have a low resolution (Neumann, 2002). Therefore, signaling behaviors that do not rely on vision are essential for mate attraction in male cicadas.

This study focuses on the acoustic communication in male *Meimuna opalifera* that inhabit East Asia. Empirical studies demonstrated that the calls of these cicadas have complex acoustic traits with remarkable melody and are divided into three calling sections (Figure 2a; Hagiwara and Ogura, 1960; Nakao, 1958). Section I contains calls with gradually increasing amplitudes, Section II contains calls with periodically changing amplitudes, and Section III contains calls with gradually decreasing amplitudes (Hayashi and Saisho, 2015). In this study, we focus on Section II and refer to the calls included in the focal section as “Type I calls” because it has the longest duration and maximum amplitude among the three sections (Nakao, 1958). Moreover, the cicadas vocalize another type of calls when other cicadas vocalize nearby (Figure 2a; Hayashi and Saisho, 2015; Saisho, 2019). In this study, we refer to these calls as “Type II calls.” Type II calls are longer than Type I calls and have a wider frequency range including that of Type I calls. In addition, Type II calls are produced intermittently but are not divided into multiple sections. Hayshi and Saisho (2015) hypothesized that Type II calls disturb the Type I calls of other males through overlapping of calls. To the best of our knowledge, however, this hypothesis has not yet been experimentally validated using sufficient data.

In this study, we examined how *M. opalifera* utilize these two types of calls. First, we recorded the calls of two males placed in a laboratory room and quantified the call types, as well as the onset and end of each call. Second, we evaluated the occurrence of acoustic interactions based on the ratio and inter-call intervals of the two call types. Finally, we tested the hypothesis of mask efficiency of Type II calls by comparing empirical data with numerical simulations.

## 2. Materials and methods

### 2.1 Laboratory experiment

We recorded the calling behavior of male cicadas (*M. opalifera*) between Aug. 19th and Sep. 19th, 2019, and between Aug. 16th and Sep. 11th, 2020. In each experiment, we collected two males per sampling occasion using a bug-catching net at the breeding site at the University of Tsukuba, Ibaraki, Japan, and immediately moved them to our laboratory. In total, the vocalizations of 46 males were recorded across the collection periods. The collected cicadas were placed into cylindrical plastic cages (height: 12.5 cm, radius: 11.0 cm) that were set 150 cm apart (Figure 1). The separation distance of 150 cm was based on similar experiments that we conducted in 2018, with different inter-cicada distances between 50 cm and 300 cm used; 150 cm was the distance at which the cicadas most frequently vocalized calls. We also placed eight microphones (cx-500F, JTS) around the cages to obtain audio data with a higher signal-to-noise ratio for each male.

**Figure 1.**
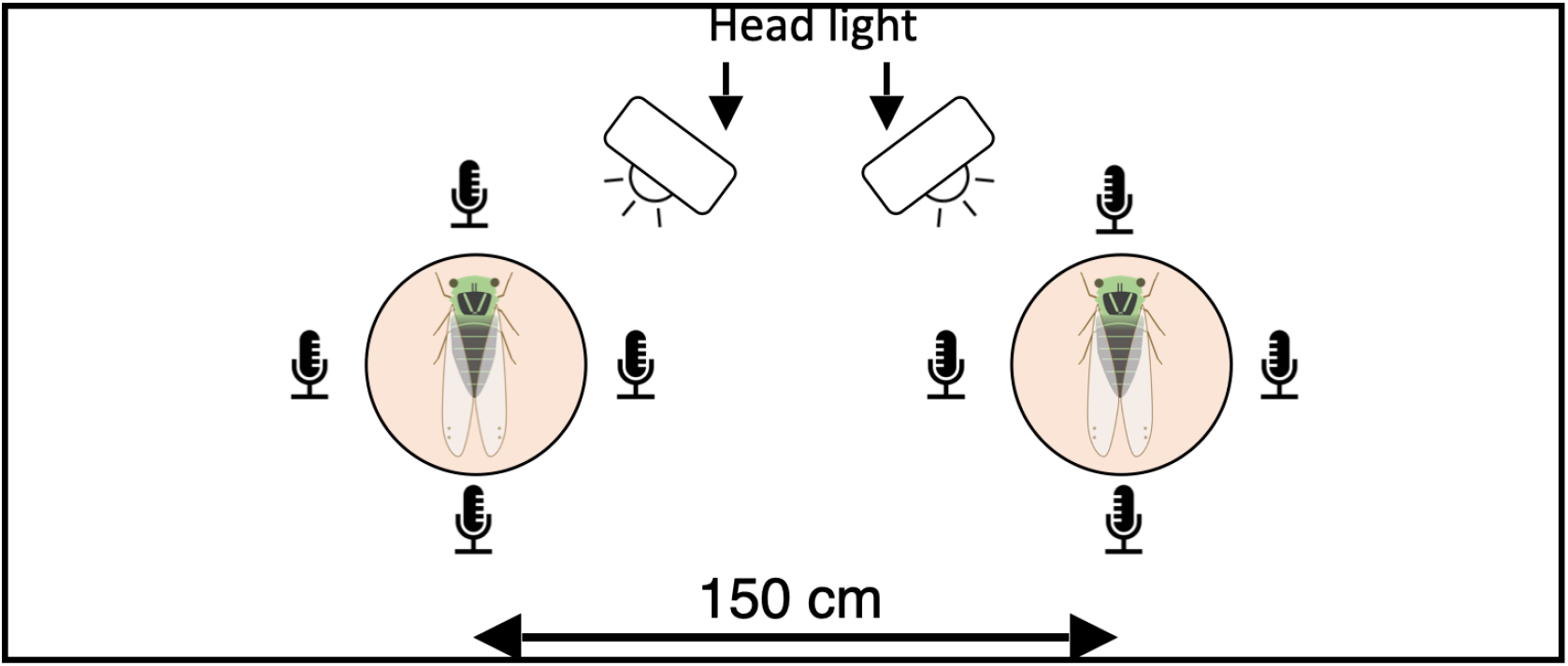
Schematic diagram describing the layout of equipment. Two male cicadas were put into cages 150 cm apart. The calls of the cicadas were recorded by eight omnidirectional microphones. The illuminance at the center of each cage was adjusted to that of natural conditions by head lights.

To elicit calling behavior in male cicadas, we adjusted for the two environmental conditions. The first condition was the illuminance of the cage. Cicadas vocalize calls depending on illuminance (Hayashi and Saisho, 2015). Therefore, we measured illuminance using an illuminance meter (SEKONIC i-346) at the breeding site and adjusted the illuminance of the cage to a value similar to natural conditions (between 2000 lx and 3000 lx) using LED lights (Tinzzi LED head light (ASIN: B0781GZCCJ) in 2019 and montbell 1124431 in 2020). The second condition was the broadcast of conspecific calls. Cicadas are known to call together in the wild (Williams and Smith, 1991), indicating that the calls of a cicada can stimulate the calling behavior of other cicadas. Therefore, we broadcast cicada’s calls recorded in a previous study (Hayashi and Saisho, 2015) for a few minutes prior to each experiment.

The calls of the cicadas were recorded using an audio recorder (R44, Roland) at 48.0 kHz sampling frequency and 16-bit quantization. Each experiment started immediately after placing the cicadas into the cages and continued between three and seven hours. As a result, we recorded approximately 125 h of the choruses from 50 males. Note that the experimental conditions were almost the same between 2019 and 2020, and only the reverberation of the room was slightly different because of the wall materials (a perforated board was used in 2019, and an anechoic material (SONEX, SOH-3) was used in 2020). All the experiments were performed in accordance with the guideline of the Animal Experimental Committee of University of Tsukuba.

### 2.2 Detection of calling time and type

Here, we explain how the call types were discriminated and how the onset and end of the calls were detected. Type I calls have the characteristics of a short duration and high repetition rate, whereas Type II calls have the characteristics of a longer duration and lower repetition rate. In Steps 1 and 2, we categorized the call types based on the difference in acoustic traits. In Step 3, we precisely estimated the onset and end of each call using the categorization result of Steps 1 and 2.

#### Step1: Approximate detection of each call

We estimated the peak and duration of the calls before categorizing call types. The procedure for this step was (1) intensity detection, (2) smoothing, and (3) normalization.

First, we calculated the intensity of the calls as the square of the audio signal.

Given the audio signal recorded by the microphone (*s*(*i*)), the intensity (*r*(*i*)) was calculated as follows:

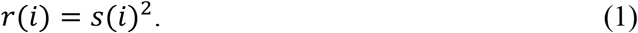

where *i* represents a specific sampling point of the data. In the experiments, we deployed four microphones for each male (Figure 1). To emphasize the signal of the target male, we squared the signals of the four nearby microphones according to Equation (1) and added them.

Next, we calculated the moving average of the squared signal (*r*(*i*)) to reduce the noise and obtain the data trend (Silvis-Cividjian, 2017; Tsokos, 2010). First, we set the sampling period with *n* data points (corresponding to the period of *n*/48.0kHz (sec)) around a specific sampling point *i*. Second, we calculated the average of the sampling period and treated it as the moving average (*M*(*i*)) at the point *i* as follows:

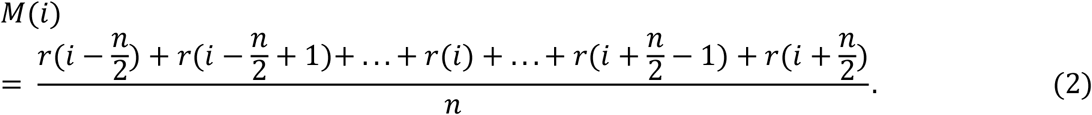

In this step, we set *n* = 24,000 (corresponding to a sampling period of 0.5 s) to detect both Type I and Type II calls.

To correct the variation in the call intensity originating from the difference in call types as well as the individuality of cicadas, we normalized the smoothed data *M*(*i*) by restricting its range between zero and one, as follows:

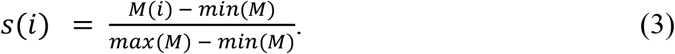

Specifically, we divided each calling bout into 200 subsections, corrected the duration of each subsection so as to start and end with a silent period and include one or more calls, and then applied Eq. (3) to each subsection. Note that division into the subsections was necessary because the focal cicada sometimes moved, causing the amplitude of the recorded audio data to change intermittently.

We detected the peak and duration of each call from the smoothed and normalized data *s*(*i*). In this step, we used the function *findpeaks* in MATLAB with an amplitude threshold of 0.5. Subsequently we successfully detected the peak and duration of most calls; however, only a few calls were missed. Because this step was only for the labeling purpose and the negative effect was minimal (see Figure S1 of Supplementary Information for details; more precise detection was conducted in Step 3), we proceeded to the next step.

#### Step2: Labeling of call types

We labeled the call types based on the acoustic characteristics calculated in Step 1. As explained in the Introduction, Type I calls have a short duration and are repeated at a higher rate in each bout. In contrast, Type II calls have a longer duration and are repeated at a lower rate. Given these differences, we applied the k-means++ method (David and Vassilvitskii, 2007) to the acoustic characteristics (i.e., the average duration and number of calls in each bout). Prior to the analysis, we normalized the ranges of the acoustic characteristics between zero and one because the k-means++ method is based on the distance between data (Silvis-Cividjian, 2017; Tsokos, 2010) and the performance can be deteriorated by the difference in the ranges of input values. Consequently, we succeeded in automatically identifying the call types (Figure 2b; note that we also confirmed the validity of the labeling by manually checking the result with the original audio data).

**Figure 2.**
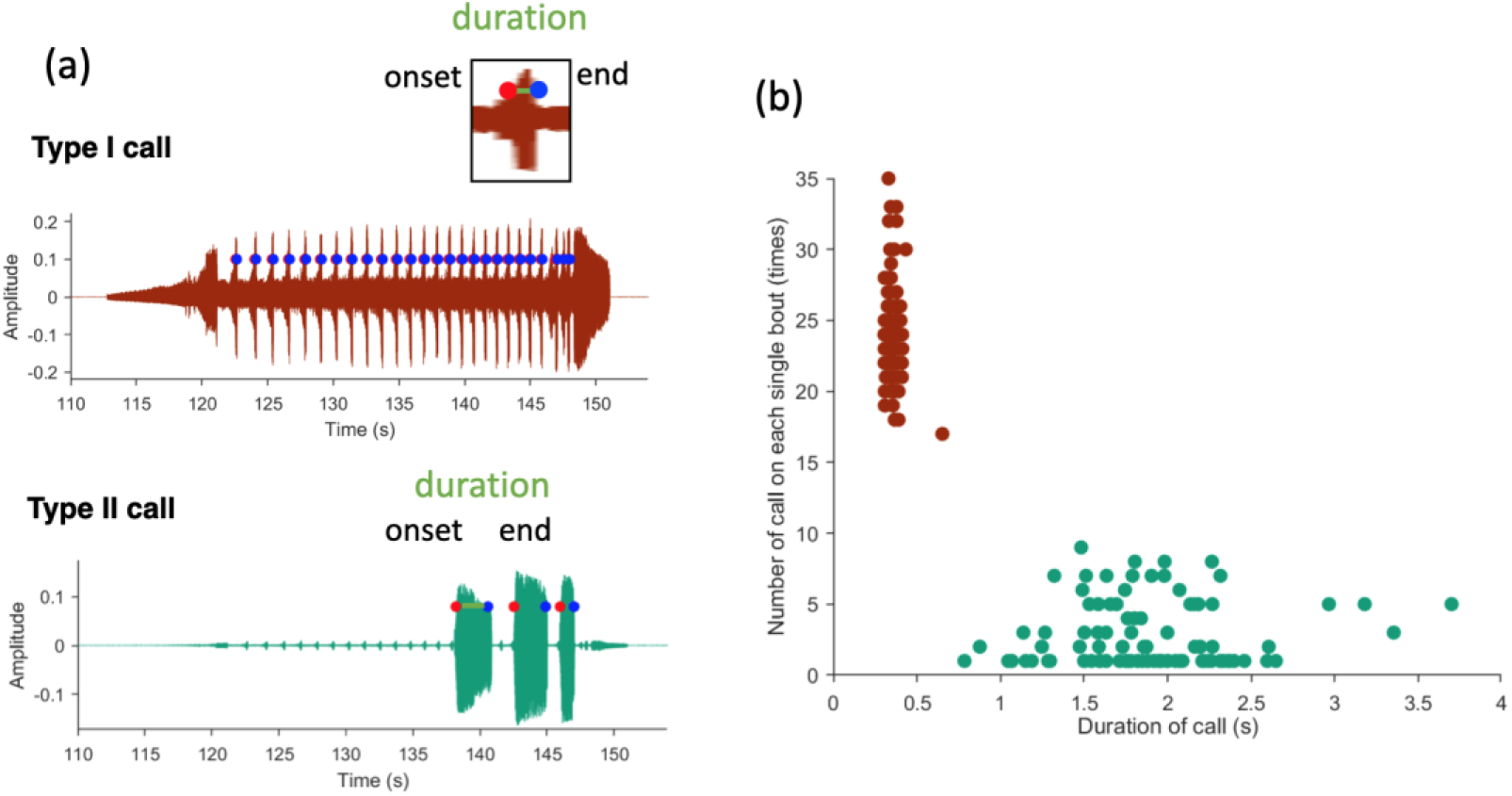
Detection and categorization of multiple call types. (a) Waveforms of Type I and Type II calls. The onset and end of each call are indicated by red and blue points, respectively. Males of this species intermittently produce successive sounds with Type I and Type II calls. We refer to a sequence of calls as “a single bout” and analyze the relationship between the two call types within each bout. (b) Categorization of Type I and Type II calls by the k-means++ method. Red and green points represent different categories obtained from the analysis, corresponding to Type I and Type II calls, respectively.

#### Step3: Accurate estimation of call traits

We estimated the onset and end of calls more accurately based on the categorization result of Step 2. Specifically, we retuned the sampling points of the moving average at 1,200 points for Type I calls (corresponding to 0.025 s) and 14,400 points for Type II calls (corresponding to 0.300 s) in order to accurately follow the trend of the respective call types. We further retuned the amplitude threshold for the call detection at the half of each call (i.e., the onset of each call was detected when the waveform *s*(*i*) first exceeded half the amplitude of the peak; the end of each call was detected when the waveform *s*(*i*) first fell below half the value of the peak). This additional procedure allowed us to estimate the onset and end of calls more accurately than in Step 1 (Figure S1 in Supplementary Information). However, only the last Type I call in each bout was relatively close to the following calling section (Figure 2a), which sometimes caused misestimations. Namely, the beginning of the following section (corresponding to Section III explained in the Introduction) was sometimes included in the data extracted for the analysis and was accordingly misestimated as a Type I call. To avoid this issue, we excluded only the last Type I call of each calling bout from the analysis. Consequently, we obtained three acoustic traits (onset, end, and duration) for each call.

### 2.3 Phase difference between Type I and Type II calls

We quantified how male *M. opalifera* responded to the calls of another male. In particular, male *M. opalifera* tend to produce Type II calls when responding to Type I calls of another male (Saisho, 2019). To quantify this timing strategy in the acoustic communication, we calculated the phase difference φ as follows (Aihara et al., 2011, Aihara et al., 2014; Pikovsky et al., 2002):

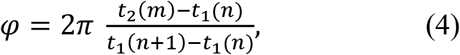

where

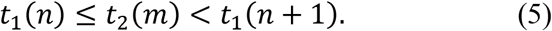

Here, *t*_1_ and *t*_2_ represent the onsets of Type I and II calls, respectively (see Figure 3a). Equation (5) indicates that the onset of the Type II call (*t*_2_(*m*)) between *t*_1_(*n*) and *t*_1_(*n* + 1) is used for the calculation, constraining the range of the phase difference from 0 to 2π. Subsequently, the phase difference of 0 or 2π indicates that a male cicada produces a Type II call at the same time as another cicada produces a Type I call. In contrast, the phase difference π indicates that a cicada produces a Type II call in the middle of the interval of Type I calls produced by another cicada. Thus, the phase difference allows us to quantify the temporal chorus structure based on its value (Aihara et al., 2011; Aihara et al., 2014).

**Figure 3.**
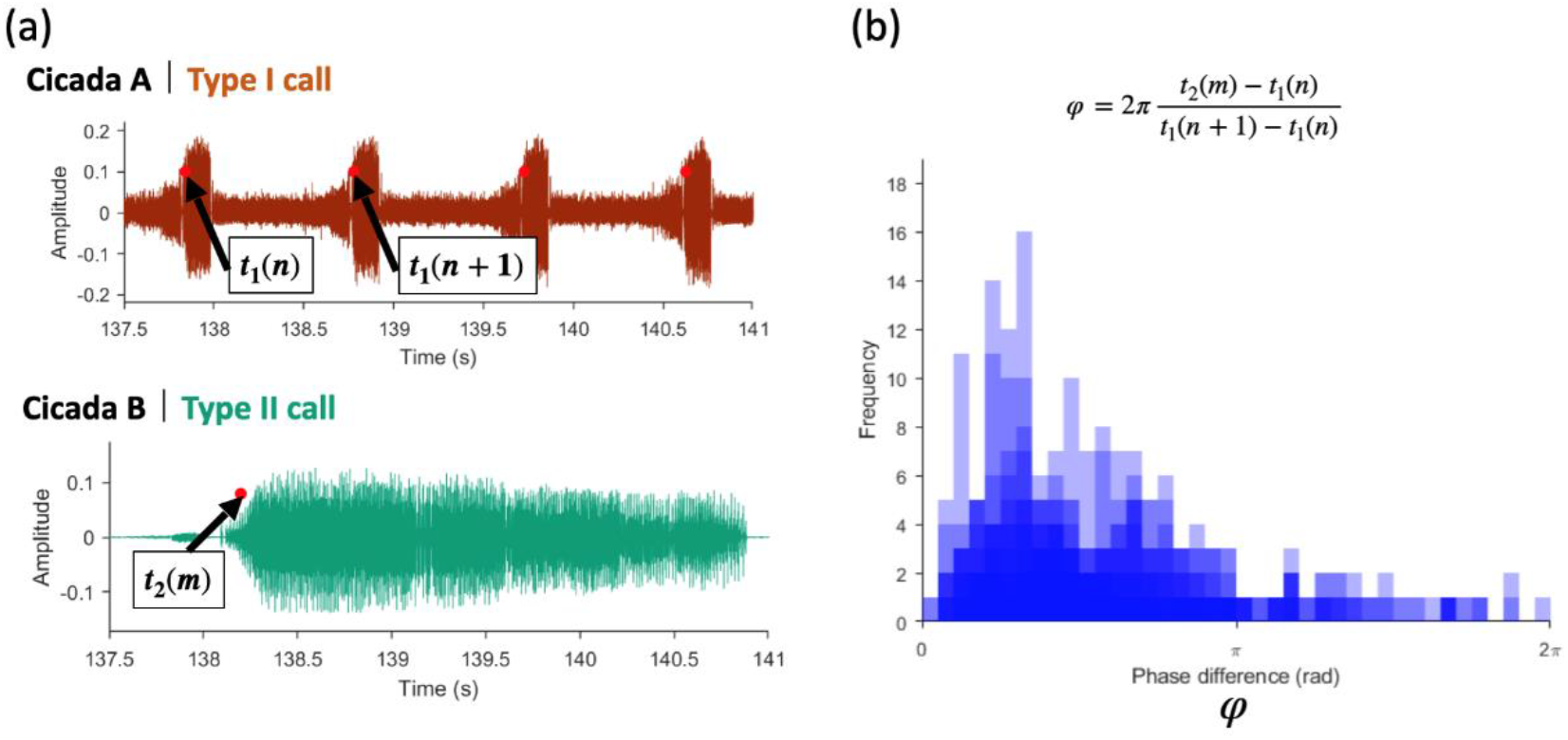
Temporal structure in chorusing bouts of male cicadas. (a) Onset times of Type I and Type II calls estimated from the audio data. (b) Distribution of a phase difference (φ). The phase difference was localized between 0 and π, indicating that a male cicada produced Type II calls immediately after another male produced Type I calls.

Next, we used the Rayleigh test to examine whether the distribution of the phase difference was localized around a specific value. The Rayleigh test allows us to test the null hypothesis that the distribution of the angular data is not localized (Fisher, 1993) (namely, the null hypothesis is rejected when the angular data is localized). In this analysis, we treated the phase differences as angular data, set the significance level to 0.001, and applied the Rayleigh test to the sequence of the phase differences using the circular package (version 0. 4. 95) in R language (version 4. 1. 3).

### 2.4 Numerical simulation of the masking performance of Type II calls

By combining empirical data and numerical simulation, we tested the hypothesis that a male *M. opalifera* vocalizes Type II calls to effectively mask the Type I calls of another male. In the simulation, we randomized the onset times of Type II calls and calculated the number of actual Type I calls masked by the randomized Type II calls for comparison with the empirical data. First, we generated a uniform random number within the calling bout of actual Type I calls of one cicada and treated it as the onset time of the randomized Type II calls of another cicada. When the generated random time partially overlapped with one of the previously generated times, we calculated the random time again until it did not overlap. Second, a Type I call was determined to be masked by a Type II call when the onset time of the Type I call was included within the duration of the Type II call. Third, we calculated the number of Type I calls masked by the randomized Type II calls in each calling bout. To reduce the uncertainty of the simulation, we generated the random onset times of Type II calls for 10,000 patterns by changing the random seed value and repeated the above procedure.

Next, we compared the number of masked calls obtained from the simulation with empirical data. When the number of masked calls in the empirical data was significantly larger than that of the simulation, we determined that the actual male cicadas effectively masked the calls of their competitors. Specifically, we set the significance level at 2.5 % (i.e., the difference was determined to be significant when the amount of the empirical data was higher or lower than the 2.5% distribution of the number of the simulations), which corresponds to a two-sided test with a significance level of 0.05.

## 3. Results

### 3. 1 Fundamental acoustic traits of each call type

Here, we summarize the fundamental acoustic traits of each call type. From the laboratory experiments, Type I and II calls were recorded 19372 and 949 times, respectively. Concerning Type I calls, the duration of the calls was 0.147 ± 0.080 s (average ± standard deviation), the number of calls included in each calling bout was 26.7 ± 4.1. For Type II calls, the duration of the calls was 2.37 ± 0.87 s, and the number of calls included in each calling bout was 2.82 ± 1.8. Thus, the duration of Type I calls was significantly shorter than that of Type II calls (19372 Type I calls, 949 Type II calls, Z = 6,608, p < 0.001 (Brunner-Munzel test with R)) and the number of Type I calls is larger than that of Type II calls (726 bouts of Type I calls, 336 bouts of Type II calls, Z = −349, p < 0.001 (Brunner-Munzel test with R)).

### 3.2 Combinations of calls

Male cicadas chose Type I calls or Type II calls for each calling bout. Here, we summarize the frequency of usage of the two call types (Figure 4a). The items in the figure are Type I and Type I; Type I and Type II; Type II and Type II; Type I and Silence; and Type II and Silence. The most common combination was Type I and Silence (382 times), followed by Type I and Type II (340 times). Type I and Type I was rarely observed (10 times). In contrast, Type II with Type II or Silence was never observed. The frequent observation of Type I and Type II is consistent with that of a previous study (Saisho, 2019).

**Figure 4.**
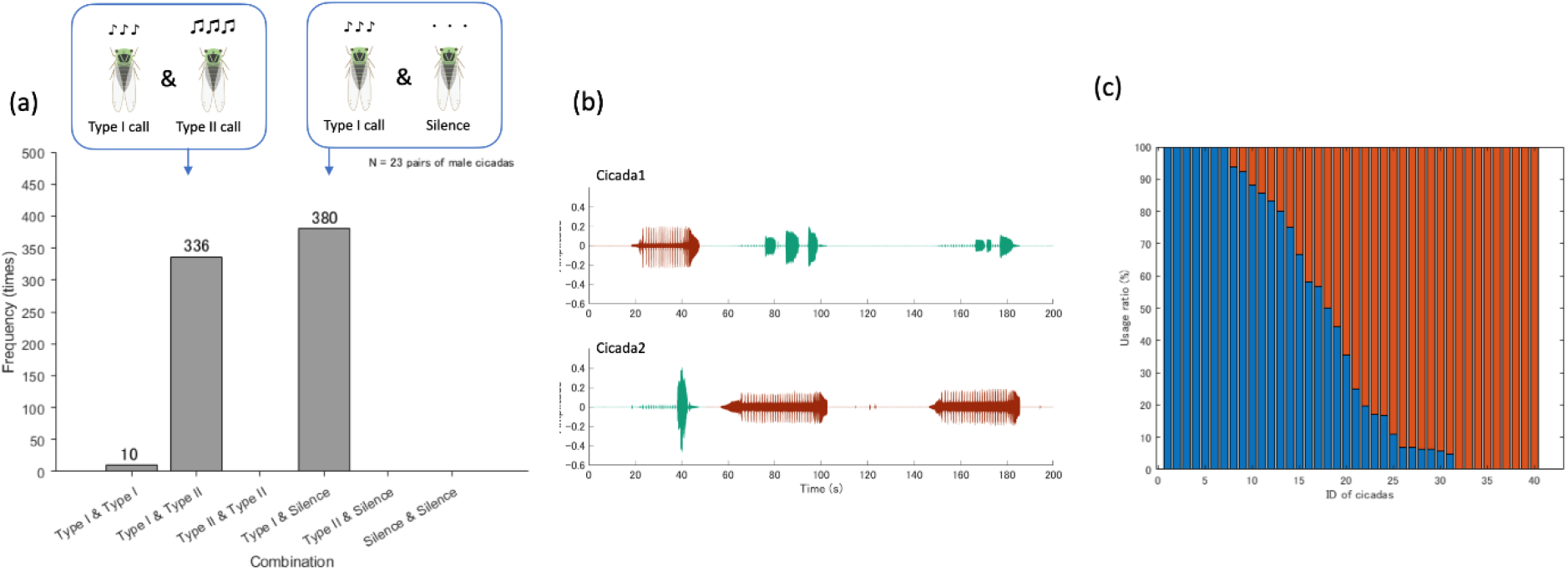
Dependence of call types on individual males. (a) The frequency of Type I calls and Type II calls for each pair of male cicadas. The abscissa shows the combination of call types. (b) Representative data showing the switching of the two call types. Initially, Cicada 1 vocalized Type I calls while Cicada 2 vocalized Type II calls. Subsequently, they switched Type I and II calls. (c) The ratio of call types for individual males. Red bars represent the ratio of the use of Type I calls, and blue bars represent that of Type II calls.

We observed switching behavior in which one cicada vocalized two call types during one experiment. Figure 4b shows the representative switching dynamics: (1) Cicada 2 first vocalized Type II calls while Cicada 1 vocalized Type I calls, and then (2) Cicada 2 switched to Type I calls whereas Cicada 1 switched to Type II calls. The ratio of the use of the two call types depended on the individual males (Figure 4c).

### 3.3 Phase difference

Figure 3b shows the distribution of the phase differences, in which the distribution of each dataset is displayed with translucent blue bars. The Rayleigh test demonstrated that the distribution was significantly different from a uniform distribution (N = 901, Z=0.5854, p < 0.001). Furthermore, the phase differences were localized around 0 to π and not around π to 2π, indicating that the cicadas vocalized Type II calls immediately after Type I calls.

### 3.4 Numerical simulation of the masking hypothesis

To test the hypothesis that male *M. opalifera* produce Type II calls to effectively mask the Type I calls of their competitors, we compared the number of masked calls of the empirical data with that of the numerical simulation. Table 1 summarizes the results for the 31 males that produced Type II calls during the experiment. Our results revealed that (1) 11 males (35.5%) did not effectively mask the calls, (2) the results were not significant for 20 males (64.5%), and (3) no male effectively masked the calls. Figure 5 shows the representative results of the empirical data (red line) and the numerical simulation (histogram with gray bars) for each case.

**Table 1.** Comparison of the number of masked calls between the empirical data and numerical simulation to indicate masking effectiveness. This result was obtained from 31 males.

|  | (1)<br>lower 2.5 %<br>(not effective) | (2) between<br>2.5 and 97.5 %<br>(not significant) | (3)<br>upper 2.5 %<br>(effective) |
| --- | --- | --- | --- |
| Frequency | 11 | 20 | 0 |

**Figure 5.**
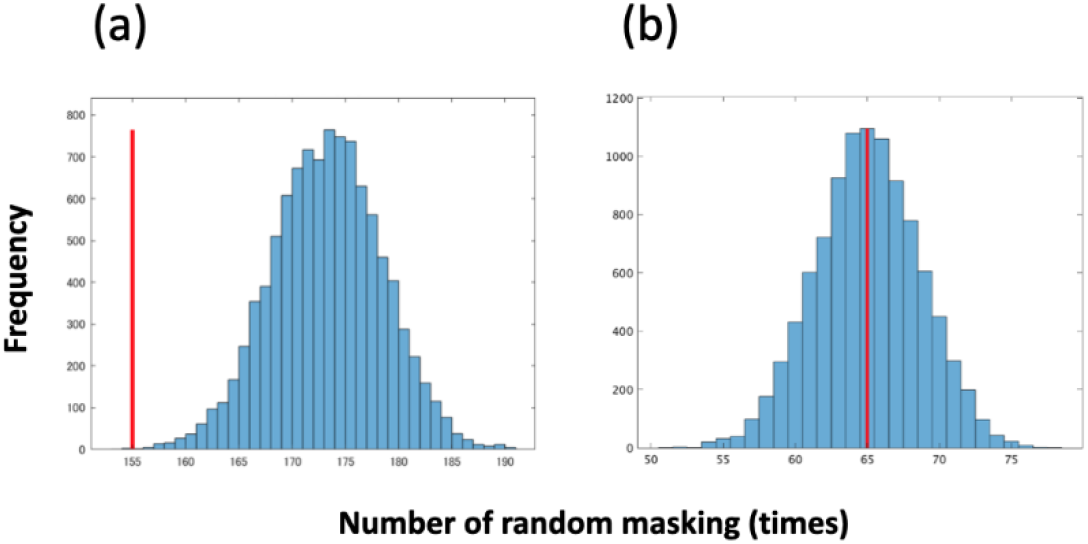
Representative results for the test of the hypothesis that male *M. opalifera* produced Type II calls to effectively mask the Type I calls of their competitor. The red line shows the number of masked Type I calls of the empirical data and the histogram shows that of the simulation. In the simulation, we estimated the number of the masked calls by randomizing Type II calls in 10,000 patterns; therefore, the number was obtained as a distribution. Each graph shows the representative results for (a) the lower 2.5% case (not effective) and (b) the case between 2.5 and 97.5% (not significant).

## 4. Discussion

We investigated acoustic communication between male cicadas (*M. opalifera*) based on two call types. The laboratory experiments demonstrated that (1) cicadas used two call types depending on the situation, and (2) they vocalized Type II calls against Type I calls at a specific short interval (a phase difference slightly larger than zero). To test the hypothesis that the timing strategy allowed the males to effectively mask the calls of their competitors, we performed numerical simulations randomizing the onset of Type II calls and compared their masking performance with the empirical data. Consequently, our hypothesis was not supported.

Given that the cicadas produced Type II calls against the Type I calls of their competitor at a specific interval, the cicadas obviously interacted through sounds when using the two call types. We speculate that the timing strategy has functions other rather than masking, such as the following:

➢ **Competition hypothesis:** The first hypothesis is that cicadas produce Type II calls to reduce the mate attraction performance of competitors or to make competitors to fly away. Because our experiments demonstrated that Type II calls masked one or more Type I calls, Type II calls may deteriorate the original effects of the Type I calls of the competitors. In both cases, male cicadas that vocalize Type II calls are likely to have an advantage in mating.
➢ **Sneaking hypothesis:** The second hypothesis states that Type II calls imitate the last long call in a melodic calling bout. Male cicadas periodically changed the amplitude of their calls over Section II (corresponding to Type I calls) and then gradually decreased the amplitude in Section III (Figure 2a; see also Introduction). In our experience, the acoustic traits (e.g., duration and frequency) of Type II calls are similar to those of Section III. If the transition from Section II to Section III is the key to mate attraction, the male producing Type II calls has an advantage in mating: namely, if a male produced Type II calls after his competitor had successively produced Type I calls, he could imitate Section III and likely attract females. Combined with the fact that the total duration of Type II calls in a calling bout is much shorter than that of Type I calls, males producing Type II calls may increase the probability of mating while reducing energy consumption.
➢ **Cooperation hypothesis:** The third hypothesis states that cicadas attempt to increase the total sound intensity of aggregation by vocalizing Type II calls. As mentioned in Section 3.2, cicadas vocalized Type II calls while another cicada vocalized Type I calls. We speculate that such an overlap of calling periods produces a louder sound volume as a whole aggregation, attracting more females overall.

To test these hypotheses, video recordings of the spatial movement of females and males are required. Given the variance in the calling behavior (Figure 4), more controllable procedures such as playback experiments using loudspeakers (Kodama and Kasuya 2022) seem more appropriate than our approach using only wild cicadas.

Next, we address the limitations of our study. In our experiments, we fixed the distance between cicadas at 150 cm and observed that cicadas frequently vocalized Type II calls against the Type I calls of their competitor (Figure 4a) and rarely vocalized Type I calls against Type I calls. While male *M. opalifera* are known to vocalize Type II calls at such short distances (Saisho, 2019), they frequently produced Type I calls at field sites where the inter-male distance is much longer than that in our laboratory experiments. Thus, the ratio of Type I and Type II calls may change depending on the distance between the males.

## Acknowledgements

We thank Dr. Ryu Takeda, Dr. Tohru Kawabe and Dr. Yoshito Hirata for their valuable comments regarding this study. We thank Mr. Masahiro Shirasaka for the discussions and assistance in building the recording equipment. We are also grateful to Mr. Makoto Nashiki, who helped with the data analysis (estimation of call timing).

## Competing interests

No competing interests declared.

## Funding

This study was partially supported by the JSPS Grant-in-Aid for Scientific Research (B) (No. 20H04144).

## Notes

### Competing Interest Statement

The authors have declared no competing interest.

## References

Aihara, I., Mizumoto, T., Otsuka, T., Awano, H., Nagira, K., Okuno, H. G. and Aihara, K. (2014). Spatio-temporal dynamics in collective frog choruses examined by mathematical modeling and field observations. Sci. Rep. 4, 1–8.

Aihara, I., Takeda, R., Mizumoto, T., Otsuka, T., Takahashi, T., Okuno, H. G. and Aihara, K. (2011). Complex and transitive synchronization in a frustrated system of calling frogs. Phys. Rev. E 83, 031913.

Arthur, D. and Vassilvitskii, S. (2007). k-means++: The Advantages of Careful Seeding. SODA ‘07 Proceedings of the Eighteenth Annual ACM-SIAM Symposium on Discrete Algorithms, pp. 1027–1035.

Au, W. W. L. and Herzing, D.L. (2003). Echolocation signals of wild Atlantic spotted dolphin (Stenella frontalis). J. Acoust. Soc. Am. 113, 598–604.

Brush, J.S. and Narins, P. M. (1989). Chorus dynamics of a neotropical amphibian assemblage: comparison of computer simulation and natural behaviour. Anim. Behav. 371, 3344.

Catchpole, C. K. and Slater, P. J. B. (2008). Bird song biological themes and variations. Cambridge, UK: Cambridge University Press.

Demartsev, V., Strandburg-Peshkin, A., Ruffner, M. and Manser, M. (2018). Vocal turn-taking in meerkat group calling sessions. Curr. Biol. 28, 3661–3666.

Fisher, N. I. (1993). Statistical analysis of circular data. Cambridge, UK: Cambridge University Press.

Fonseca, P. J. (2014) Cicada Acoustic Communication. In: Hedwig B. (eds) Insect Hearing and Acoustic Communication. Animal Signals and Communication, vol 1. Heidelberg, Germany: Springer Berlin.

Fujioka, E., Aihara, I., Sumiya, M., Aihara, K. and Hiryu, S. (2016). Echolocating bats use future target information for optimal foraging. Proc. Natl. Acad. Sci. USA 113, 4848–4852

Gerhardt, H. C. and Huber, F. (2002). Acoustic communication in insects and anurans. Chicago, USA: University of Chicago Press.

Greenfield, M. D. and Roizen, I. (1993). Katydid synchronous chorusing is an evolutionarily stable outcome of female choice. Nature 364, 618–620.

Greenfield, M. D., Aihara, I., Amichay, G., Anichini, M. and Nityananda, V. (2021). Rhythm interaction in animal groups: selective attention in communication networks. Philos. Trans. R. Soc. B 376(1835), 20200338.

Hagiwara, S. and Ogura, K. (1960). Analysis of songs of Japanese cicadas. J. Insect Physiol. 5, 259–263.

Hayashi, M. and Saisho, Y. (2015). The Cicadidae of Japan. Seibundo-shinkosha. (in Japanese)

Jones, D. L., Jones, R. L. and Ratnam, R. (2014). Calling dynamics and call synchronization in a local group of unison bout callers. J. Comp. Physiol. A 2001, 93107.

Kodama, T. and Kasuya, E. (2022). The song of the cicada Meimuna opalifera; do the functions differ between the former and the latter part of the song? ESJ69, Abstract, Poster presentation 1–110.

Nakao, S. (1958). On the diurnal variation of a song-length of Meimuna opalifera Walker and the effect of population density upon it. Japanese Journal of Entomology. 26(4) 201–209 (In Japanese, only summary in English).

Neumann, T. R. (2002). Modeling Insect Compound Eyes: Space-Variant Spherical Vision. In International Workshop on Biologically Motivated Computer Vision, pp. 360–367.

Ota, K., Aihara, I. and Aoyagi, T. (2020). Interaction mechanisms quantified from dynamical features of frog choruses. R. Soc. Open Sci. 7, 191693.

Pikovsky, A., Rosenblum, M. and Kurths, J. (2002). Synchronization: a universal concept in nonlinear science. Cambridge University Press, Cambridge.

Rhinehart, T. A., Chronister, L. M. and Devlin, T. (2020). Acoustic localization of terrestrial wildlife: current practices and future opportunities. Ecol. Evol. 10, 1–25.

Saisho, Y. (2019). The Handbook of Cicadas. Bun-ichi Co.,Ltd. (In Japanese).

Silvis-Cividjian, N. (2017). Sound Processing. Pervasive Computing. Springer, pp. 86–87.

Suzuki, R., Matsubayashi, S., Saito, F., Murate, T., Masuda, T., Yamamoto, K., Kojima, R., Nakadai, K. and Okuno, H. G. (2018). A spatiotemporal analysis of acoustic interactions between great reed warblers (Acrocephalus arundinaceus) using microphone arrays and robot audition software HARK. Ecol. Evol. 8, 812–825.

Takahashi, D. Y., Narayanan, D. Z. and Ghazanfar, A. A. (2013). Coupled oscillator dynamics of vocal turn-taking in monkeys. Curr. Biol. 23, 2162–2168 (doi:10.1016/j.cub.2013.09.005).

Tsokos, C. P. (2010). K-th moving, weighted and exponential moving average for time series forecasting models. Eur. J. Pure Appl. Math. 3(3), 406–416.

Wells, K. D. (2007). The ecology and behavior of amphibians. Chicago, USA: The University of Chicago Press.

Willi, A. R. and Zeil, J. (2015). The visual system of the Australian ‘Redeye’ cicada (Psaltoda moerens). Arthropod Struct. Dev. 44(6), Part A, 574–586.

Williams, K. S. and Smith, K. G. (1991). Dynamics of Periodical Cicada Chorus Centers (Homoptera: Cicadidae: Magicicada). J. Insect Behav. 4(3), 275–291.

Young, D. and Bennet-Clark, H. C. (1995). The role of the tymbal in cicada sound production. J. Exp. Biol. 198(4), 1001–1020.

