## Supplemental Figure 1 for "Temporal structure of two call types produced by competing male cicadas"

**Supplementary information**

**Additional procedure at Step 3**

Here, we explain how to tune the sampling rate of the moving average and the procedure to correct the misdetection of Type I calls. For the precise detection of Type I calls, we need to set a suitable value of the sampling rate for the calculation of the moving average. Accordingly, we increased the sampling rate from 24,000 points (corresponding to 0.5 s) to 1,200 points (corresponding to 0.025 s) for the detection of Type I calls and increased the rate from 24,000 points (corresponding to 0.5 s) to 14,400 (corresponding to 0.3 s) for the detection of Type II calls as explained in Step 3. Figure S1 demonstrates that this procedure allowed us to more precisely follow the trend of each call type. However, the misdetection of a few calls occurred at the boundary between Section II (the section including Type I calls) and Section III of the melodic calling bout (see Introduction for details). This is because the estimated calling bouts sometimes included just the beginning of Section III. Accordingly we sometimes misestimated the beginning of Section III as a Type I call. To avoid this misdetection, we excluded the last Type I call in each bout from the analysis.

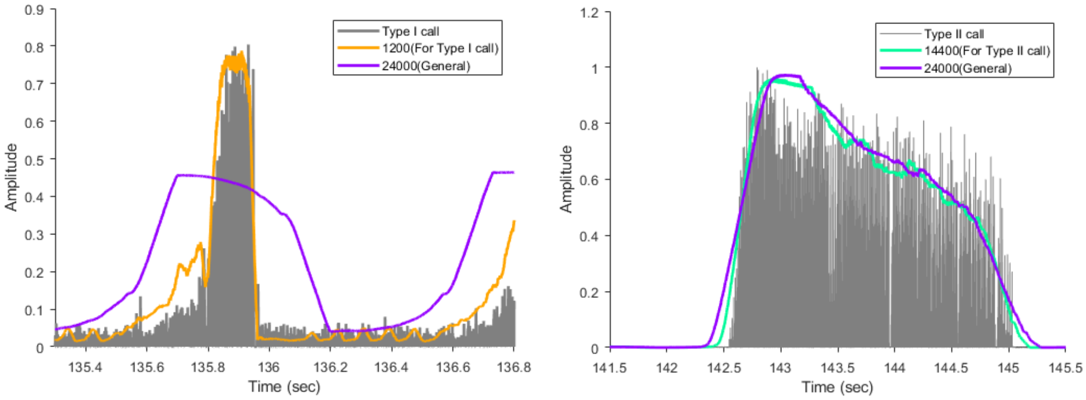


Figure S1. The sampling rates for the detection of each call type: (A) Type I calls and (B) Type II calls. The increase of the sampling rate allows us to more precisely follow the trend of each call type.
